# Rat perichondrium transplanted to articular cartilage defects forms articular-like, hyaline cartilage

**DOI:** 10.1101/2020.11.27.401091

**Authors:** Zelong Dou, Daniel Muder, Marta Baroncelli, Ameya Bendre, Alexandra Gkourogianni, Lars Ottosson, Torbjörn Vedung, Ola Nilsson

## Abstract

Reconstruction of articular surfaces destroyed by infection or trauma is hampered by the lack of suitable graft tissues. Perichondrium autotransplants have been used for this purpose. However, the role of the transplanted perichondrium in the healing of resurfaced joints have not been investigated. Perichondrial and periosteal tissues were harvested from rats hemizygous for a ubiquitously expressed enhanced green fluorescent protein (EGFP) transgene and transplanted into full-thickness articular cartilage defects at the trochlear groove of distal femur in wild-type littermates. As an additional control, cartilage defects were left without a transplant (no transplant control). Distal femurs were collected 3, 14, 56, 112 days after surgery. Transplanted cells and their progenies were readily detected in the defects of perichondrium and periosteum transplanted animals but not in defects left without a transplant. Perichondrium transplants expressed SOX9 and with time differentiated into a hyaline cartilage that expanded and filled out the defects with *Col2a1-*positive chondrocytes and a matrix rich in proteoglycans. Interestingly, at later timepoints the cartilaginous perichondrium transplants were actively remodeled into bone at the transplant-bone interface and at post-surgery day 112 EGFP-positive perichondrium cells at the articular surface were positive for *Prg4*. In addition, both perichondrium and periosteum transplants contributed cells to the subchondral bone and bone marrow, suggesting differentiation into osteoblast/osteocytes as well as bone marrow cells. In summary, we found that perichondrium transplanted to articular cartilage defects develops into an articular-like, hyaline cartilage that integrates with the subchondral bone, and is maintained for an extended time. The findings indicate that perichondrium is a suitable tissue for repair and engineering of articular cartilage.

## Introduction

During development, bone formation is initiated by the condensation of immature mesenchymal cells that eventually lead to bone formation either by the intramembranous or endochondral route. In intramembranous bone formation a differentiation program controlled by transcription factors Runx2 and Osx, Wnt signaling and other factors leads to direct differentiation of preosteoblast and subsequently osteoblasts and osteocytes, which lay down a matrix rich in collagen type I (Col1) that is mineralized. In endochondral bone formation, mesenchymal cells undergo a program controlled by the chondrogenic transcription factor SOX9 and differentiate into chondrocytes that proliferate and produce a matrix rich in collagen type II (Col2) and proteoglycans forming the initial cartilaginous template in which chondrocytes will undergo hypertrophic differentiation switching from Col2 to collagen type X (ColX) expression, mineralize their matrix, and produce factors, e.g. vascular endothelial growth factor (VEGF) that attract invading vessel and bone cells that remodel the template into bone^(1,2)^. The cartilage templates are surrounded by perichondrium whereas bone tissue is surrounded by periosteum. Perichondrium and periosteum are similar in that they both consists of an outer fibrous layer and an inner cambium layer containing mesenchymal stem cells with both chondrogenic and osteogenic properties ^(3,4)^. However, they have different functions during skeletal development, maintenance and repair ^(5-7)^.

Joint formation occurs concomitantly with skeletal development and is first visible with the formation of interzones constituted of elongated, densely packed cells characterized by expression of GDF5, Gli3, Wnt4 and Wnt9a ^(8-10)^, and low expression of Matn1 and Col2 ^(11)^ and will eventually give rise to much of the synovial joint structures including the articular cartilage ^(8,12)^. As the future joint cavitates, interzone cells closest to the cavity will start producing lubricin *(Prg4)* ^(12,13)^ and these *Prg4* positive cells will serve as progenitors for the underlying layers of the articular cartilage during postnatal life ^(14,15)^. The synovial membrane produces synovial fluid, which is a blood-ultrafiltrate rich in hyaluronic acid, lubricin, and proteases, that fills up the joint cavity, and act together with lubricin expression at the joint surfaces to minimize friction. Low friction together with normal myogenesis and movement is crucial to the development of the mature joint ^(16)^.

Articular cartilage injury due to trauma, infection or degenerative diseases are common causes of suffering and disability. These injuries can occur in all joints and appear throughout all ages with massive socioeconomic consequences. Moreover, these injuries are difficult to treat. One key issue is the poor regenerative capacity of articular cartilage ^(17,18)^. Although healing may occur under certain biological conditions, it is often absent or incomplete ^(18)^. The reasons for this inability to regenerate is not clear, but likely due to the structural and physiological nature of this avascular and highly specialized tissue containing a limited number of chondrocytes with slow turnover ^(19,20)^. Consequently, a variety of surgical techniques have been introduced in attempts to repair or regenerate articular cartilage, including Pridie drilling and microfracturing methods ^(19,21)^, autologous osteochondral transplantation (mosaicplasty) ^(22-24)^, osteochondral allografting ^(19)^, autologous chondrocyte implantation ^(25)^, periosteal transplantation ^(26)^, and perichondrial transplantation ^(27)^.

The idea of using perichondrial transplantation to reconstruct articular cartilage surfaces originally stems from early observations that perichondrium has a high potential to produce cartilaginous tissue, e.g. in “cauliflower ear”, the typical malformation that wrestlers develop after bumping and twisting their ears towards the opponent’s head. The trauma causes a hemorrhage that lifts the perichondrium from the underlying cartilage and new cartilage is formed in the blood filled space ^(28)^. This and subsequent observations that not only blood, but also the synovial joint microenvironment was able to induce a chondrogenic program in the perichondrium led to the use of free perichondrium transplants to repair and resurface injured joints ^(27,29-31)^. The method has been used in a variety of patients and joints, often with variable results ^(32-35)^. Due to the need for a second surgical site at the ribcage, together with variability in the reported outcomes as well as improved orthopedic implants, the use of perichondrium transplantation declined. However, the technique is still in use and follow-up of patients treated with perichondrium transplants has indicated that the technique can produce functional articular surfaces that last over extended periods ^(36)^, thus suggesting that perichondrium has the capacity to form an articular cartilage-like surface. However, the contribution of the transplanted perichondrium to the new joint surface and the quality of the resulting articular surfaces have been discussed^(21)^ but not formally investigated. In order to address these questions, we used an inbred strain of rats with ubiquitous expression of enhanced green fluorescent protein (EGFP) from which perichondrium and periosteum were harvested and transplanted into engineered articular cartilage injuries in EGFP-negative littermates, thus allowing for cell-tracing of transplanted cells and their progenies and evaluation of their contribution to and the quality of the resulting articular surfaces during different phases of healing.

## Materials and methods

### Animals

All animal procedures were approved by the regional animal ethics committee in Stockholm (Permit no. N248/15, N118/16, and 15635-2017). In order to trace transplanted cells, inbred Lewis rats overexpressing EGFP under the ubiquitous CAG promote (Lew-Tg(CAG-EGFP)YsRrrc; Rat Research and Resource Center, Columbia, MO, USA) were bred to wild-type resulting in a 1:1 ratio of transgenic hemizygous to wild-type animals. EGFP positive animals were used as donors and wild-type littermates as recipients (n = 3 for each group and time-point, except for the no transplant group at day 14 post-surgery (n = 2)). Animals were housed under standard conditions with a 12-hour light/dark cycle and received standard rat chow and fresh water ad libitum.

### Full thickness articular cartilage injury repair using perichondrial or periosteal transplants

Six-week-old *EGFP* transgenic rats were euthanized by CO_2_ inhalation and with microsurgical technique perichondrial grafts were harvested from the cartilaginous part of the sternal ribs, and periosteal grafts were harvested from the anterior medial part of the tibia. To dissect the perichondrium grafts, the ventral part of the rib cage was cut loose with scissors. Under 4.5x loupe magnification two or three of the lower sternal ribs were detached from the sternum and separated from intercostal muscles with scalpel and forceps. The rib was put in a petri dish and kept moist with a few drops of saline. By use of a dissecting microscope (Carl Zeiss Stemi DV4 Series, Carl Zeiss, Oberkochen, Germany) with 10-40x magnification the cartilaginous part was distinguished from the osseus part of the rib. Under the microscope, a longitudinal cut was made through the perichondrium only, followed by careful dissection with micro scalpel and forceps peeling off the graft from the underlying cartilage. The perichondrial transplants were inspected and any residual cartilage was carefully removed without injuring the cambium (inner) layer of the perichondrium. The periosteal grafts were peeled off the underlying medial part of the tibia in a similar fashion using the microscope. The most intact parts of the grafts were cut into suitable sizes and kept moist in saline.

*EGFP*-negative littermates were sedated by inhalation of isoflurane/oxygen in a cage and then maintained on mask isoflurane/oxygen. Heating blanket was used to maintain the body temperature of the recipient animals during the surgery. Knees were shaved and scrubbed with 70% isopropryl alcohol, petrolatum eye ointment was administered to each eye, and normal saline (5-10 ml) subcutaneously. The animals were induced with isoflurane and kept under general anesthesia using mask isoflurane. Knee joints were opened by a longitudinal incision lateral to the patella, which in turn was dislocated in medial direction, exposing the femoral condyles. At the intercondylar groove of the knee joint, 2 partly overlapping holes were punched out by using a 16-gauge Jamshidi bone marrow biopsy needle (BD Biosciences., NJ, USA). The edges of the holes were trimmed with a surgical scalpel and the bottom of the defect was curetted to expose the bleeding subchondral bone, creating an approximately 2×5 mm large full thickness articular cartilage injury. Trimmed *EGFP*-positive perichondrial or periosteal grafts were fitted to cover the defect and placed with the cambium layer facing towards the joint space. Attachment was made under the microscope with osteosutures at each corner of the transplant by using 8-0 none-absorbable monofilament. Two sutures were put in place and tied. The graft, still partially loose, was carefully lifted and a small amount of fibrin glue (TISSEEL, Baxter Healthcare Corporation, Westlake, CA, USA) was placed under the graft. Slight pressure was evenly applied to the graft for 60 seconds before completing the attachment by placing the two remaining sutures. Excess glue was carefully removed. The joint capsule and skin were closed in layers using 7.0 braded, resorbable sutures (Ethicon, Somerville, NJ, USA). OPSITE Spray (Smith & Nephew, London, United Kingdom) was used to prevent potential inflammation caused by maceration. Bupivacaine (2.5 mg/ml) was injected around the surgical scar for local anesthesia. After the completion of surgery, meloxicam (1 mg/kg) or buprenorphine (20-30 mcg/kg) was administered every 12 hours during the first 48 hours. At post-surgery day 3, 14, 56 and 112, animals received an injection of 5-bromo-2’-deoxyuridine (BrdU; 50mg/kg) and were killed 4hrs later by carbon dioxide inhalation. Distal femoral epiphyses were rapidly excised, fixed (10% formalin) overnight, decalcified (15% EDTA, 0.5% PFA) for 4 to 6 weeks, and photographed (Leica M320 dental microscope, Leica, Wetzlar, Germany) before being divided into two halves by a frontal cut in the center of the injury site and embedded in paraffin or optimal cutting tissue compound (OCT; Histolab, Västra Frölunda, Sweden) cyomount. Paraffin sections (6μm) were placed onto TruBOND 380 slides (Electron Microscopy Sciences, Hatfield, PA, USA) and used for immunohistochemistry, *in situ* hybridization and histological stainings. Frozen sections (10μm) were placed onto SUPERFROST PLUS slides (Thermo Fisher Scientific, Waltham, MA, USA) and used for immunofluorescence.

### Histological stainings

Masson’s trichrome Staining was applied according to the manufacturer’s instructions of the Trichrome Stain (Masson) Kit (HT15-1KT, Sigma-Aldrich, Heatherhouse, United Kingdom). For Safranin O/Fast green staining, briefly, paraffin embedded tissue sections were baked at 65°C for 45 min, deparaffinized in xylene, rehydrated through an ethanol series (100%, 100%, 95%, and 70%), and rinsed in DEPC-treated water, followed by staining with Weigerts Hematoxylin for 5 min, washed in distillated water, 2s of Acid Alcohol incubation, wash in distillated water, 0.02% Fast Green incubation for 5min, 1% acetic acid incubation for 15s, then by 30min of Safranin O incubation, rinse in 95% ethanol for 5min, dehydrated in an 95% 100% ethanol, cleared in xylene, and mounted with permount (Fisher Scientific, Fair Lawn, NJ, USA). Staining was visualized by scanning the slides under bright field microscopy with a Pannoramic MIDI II 2.0.5 digital scanner (3DHISTECH Ltd, Budapest, Hungary).

### Tracing of transplanted cells using GFP Immunohistochemistry

In order to detect EGFP-positive cells, paraffin embedded tissue sections were baked at 65°C for 1 hour, deparaffinized in xylene, rehydrated through an ethanol series (100%, 100%, 95%, and 70%), and rinsed in PBS. Antigen retrieval was performed using proteinase K (10 μg/ml in PBS) at room temperature for 15 min. Endogenous peroxidase activity was blocked by incubation in 3% H_2_O_2_ at room temperature for 15 min and non-specific binding was blocked by a 1h incubation in tris-buffered saline with tween 20 (0.1M Tris, 0.15M NaCl, 0.3% v/v Tween-20, pH7.5; TBST) containing 1% BSA and 10% goat serum before incubation with the GFP primary antibody (anti-GFP antibody, ab290, Abcam, Cambridge, MA) overnight at 4°C. Staining was performed using a VECTASTAIN ABC Kit (Vector Laboratories, Burlingame, CA, USA) followed by a DAB Substrate Kit (Vector Laboratories) according to the manufacturer’s instructions. All wash steps were carried out using 0.1% TBST. Tissue sections were counterstained with methyl green (Vector Laboratories), dehydrated in an ethanol series (95%, 100%, and 100%), cleared in xylene, and mounted in permount.

### Immunofluorescence

In order to detect EGFP-positive cells of the donor allografts, OCT embedded tissue sections were thawed and dried at 37°C for 30min, and rinsed in PBS 3 times to remove OCT. Antigen retrieval was performed in proteinase K (10 μg/ml in PBS) at room temperature for 3 min, followed by incubation in TBST containing 1% BSA and 10% donkey serum for 1h before incubation with primary antibody against GFP (#ab290, Abcam) or SOX9 (anti-SOX9 antibody NBP1-85551, Novus Biologicals, Littleton, CO, USA) overnight at +4°C. The conjugated secondary antibody (Alexa Fluor 488, A-21206, Thermo Fisher Scientific) was incubated according to the manufacturer’s instructions. All wash steps were performed using 0.1% TBST. Cell nuclei were stained with DAPI (D1306, Thermo Fisher Scientific), and slides mounted in ProLong™ Gold Antifade Mountant (P36930, Thermo Fisher Scientific). Confocal (ZEISS LSM 700) images were obtained under channels of GFP and DAPI for a z stack scanning of the mounted sections. Scanning raw data was collected and imported to open resource software ImageJ (NIH Image, National Institutes of Health (NIH), Bethesda, Maryland) for brightness and contrast adjustment, followed by compression of the total scanning slices to have the final display.

### In situ hybridization of chondrocyte differentiation markers

Once the transplanted graft was localized by EGFP immunohistochemistry, in situ hybridization for chondrocyte markers *Prg4, Col10a1, Col2a1* and *Col1a1* were performed on consecutive sections. The gene sequences for rat *Col1a1* (bone and chondrocyte dedifferentiation marker), *Col2a1* (chondrocyte marker) *Col10a1* (hypertrophic chondrocyte marker), and *Prg4* (superficial chondrocyte marker) were obtained from the UCSC Genome Browser. Primers were designed using Primer-Blast (https://www.ncbi.nlm.nih.gov/tools/primer-blast/), and the resulting amplicons were confirmed by NCBI Nucleotide Blast. DNA templates for riboprobe transcription were amplified by PCR using the following reagents: Platinum Taq DNA Polymerase (Invitrogen, Carlsbad, CA, USA), cDNA reverse transcribed from total RNA isolated from 3-day-old rat proximal tibial epiphyses using a previously described protocol,^(37)^ forward primers containing a T7 promoter (5’-TAATACGACTCACTATAGGGAG-3’), and reverse primers containing a Sp6 promoter (5’-TGGATTTAGGTGACACTATAGAAG-3’): Primer sequences of Rat *Col10a1 and Prg4*,^(38)^ primer sequences of *Col2a1*^(39)^ were the same as previously illustrated. For Col1a1, forward primer (5’-CATTGGTAACGTTGGTGCTCCT-3’) and reverse primer (5’-TCTCCTCTCTGACCGGGAAGA-3’) were designed based on its cDNA (2618-2968 bp of GenBank accession no. BC133728).

PCR of DNA templates was performed with a 2720 Thermal Cycler (Applied Biosystems, Waltham, MA, USA) using the following parameters: hold at 94°C for 5 min, followed by 30 cycles of denaturing at 94°C for 30 sec, annealing at 58°C for 30 sec, and extending at 72°C for 45 sec, followed by a final extension at 72°C for 3 min. PCR products were purified by agarose gel electrophoresis and a QIAquick Gel Extraction Kit (Qiagen, Hilden, Germany). A second PCR was performed using the same parameters and the products were purified with a QIAquick PCR Purification Kit (Qiagen). Single stranded riboprobes were transcribed with a Roche DIG Labeling Kit (Roche, Basel, Switzerland) incorporating a digoxigenenin (DIG)-conjugated uracil every 20 to 25 nucleotides. Sp6 polymerase was used for antisense strand riboprobes and T7 polymerase was used for sense strand riboprobes. Riboprobes were purified with Micro Bio-Spin 30 Columns (Bio-Rad, Hercules, CA, USA) and quantified with a NanoDrop Spectrophotometer (Thermo Fisher Scientific).

Non-radioactive digoxigenin *in situ* hybridization was performed as previously described with slight modifications^(40,41)^. The detailed protocol is available upon request. Briefly, tissue sections were baked at 65°C for 1 hour, deparaffinized in xylene, rehydrated through an ethanol series (100%, 100%, 95%, and 70%), and rinsed in DEPC-treated water. Tissue sections (6 μm) were permeabilized with proteinase K at room temperature for 15 min (10 μg/ml in PBS, pH7.4), postfixed for 5 min (10% formalin), and acetylated for 15 min (0.25% acetic anhydride in 0.1M triethanolamine) with each step followed by two 5 min washes in PBS. Prehybridization was carried out at 65°C for 2 hrs in hybridization solution (50% formamide, 10mM Tris pH7.6, 200 μg/ml Torula yeast RNA, 1X Denhardt’s solution, 10% dextran sulfate, 600mM NaCl, 0.25% SDS, 1mM EDTA, pH8.0). Hybridization with DIG-labeled riboprobes (100 ng in 100 μl hybridization solution) was performed at 65°C for 4hs.

Posthybridization was carried out by washing with 50% formamide in 1xSSC at 65°C for 30 min, digesting with RNAse A (20 to 2000 μg/ml in 1M NaCl, 10mM Tris HCl, 1mM EDTA, pH8) at 37°C for 30 min, and washing in SSC at increasing stringency (4x, 1x, 0.5x, and 0.2x). For detection of hybridized riboprobes, tissue sections were rinsed in MABT (0.1M maleic acid, 0.15M NaCl, 0.1% v/v Tween-20, pH7.5), blocked with 1% BSA in MABT at room temperature for 30 min, incubated with alkaline phosphatase-conjugated anti-DIG antibody (Roche) in 1% BSA in MABT at 4 degree for overnight, and then incubated with nitro blue tetrazolium chloride/5-bromo-4-chloro-3-indolyl phosphate (NBT/BCIP) substrates (Sigma-Aldrich) in NTM (100mM NaCl, 100mM Tris pH9.5, 50mM MgCl_2_) at room temperature protected from light for 0.5-4 hrs. To mount, tissue sections were rinsed in PBS for 5 min, fixed in 10% formalin for 20 min, counter stained with nuclear fast red, dehydrated in an ethanol series (70%, 95%, and 100%), cleared in xylene, and mounted with permount. Staining was visualized by scanning the slides under bright field microscopy with a Pannoramic MIDI II 2.0.5 digital scanner (3DHISTECH).

### Histomorphometry

High resolution bright-field microscopy images were obtained by using a panoramic MIDI II 2.0.5 digital scanner and analyzed by using compatible CaseViewer software (both from 3DHISTECH). Analysis was carried out in the EGFP positive graft tissues (perichondrium/periosteum). For thickness/height of graft tissues, 10 individual measurements were obtained on GFP immunofluorescence/immunohistochemistry images and then averaged. Scar-tissue thickness was similarly measured on Masson’s trichrome stained images for the no transplant control samples. In the transplant and the repair tissues, BrdU labeling and SOX9 positive cell indices were calculated by dividing the number of positive cells in the tissues by the total number of cells. Total number of nuclei in the graft tissues were counted using Image J software. Matrix proteoglycan was assessed by Safranin O staining and the positive area measured using Caseviewer (3DHISTECH). Similarly, *in-situ* hybridization images were used to quantify the percentage area positive for type I and type II collagen expression. GFP labeling index was calculated by dividing the total number of GFP labeled cells in the subchondral bone by the total number of GFP labeled cells.

### Quantitative real-time PCR of chondrocyte marker expression

Perichondrial and periosteal grafts were investigated for expression of chondrogenic markers. Briefly, after dissection the grafts were lysed in solution C (4M guanidine thiocyanate, 25mM sodium citrate pH7, 0.1M β-mercaptoethanol), and stored at −80°C until use. Total RNA (50ng) was extracted and reverse transcribed into cDNA by Superscript Reverse Transcriptase IV (Thermo Fisher Scientific) as previously described^(39)^. Expression of chondrogenic markers (*Col2a1, Col10a1, Acan, SOX9* and *Prg4*) was quantified by real-time PCR ABI Prism 7900 Fast Sequence Detector (Thermo Fisher Scientific, Waltham, MA, USA) (primer sets are listed in Supplementary Table 1). Relative expression was calculated relative to 18S rRNA (eukaryotic 18S rRNA endogenous control, Thermo Fisher Scientific) by using the formula: 2 ^−ΔCt^ *10^6^, with ΔCt being target gene expression relative to 18S.

### Statistical analyses

Statistical analysis was carried out using Prism 8.0 software (GraphPad, San Diego, CA, USA). All data are expressed as the mean ± SEM. Two-way ANOVA with post-surgery time-points and tissues (perichondrium vs. periosteum vs. no transplant) as the two independent variables followed by relevant pair-wise comparisons using Sidak’s/Tukey’s multiple comparisons tests to determine statistical significance among study groups. Statistical significance was recognized at two-sided p-value less than 0.05.

## RESULTS

### Perichondrium and periosteum graft transplantation and cell tracing

Graft recipient animals recovered rapidly after the surgery and were able to ambulate on their hind legs immediately after recovery from anesthesia and were all ambulating normally within 24 hrs including the animals in which the injuries were left without a transplant. None of the animals developed postsurgical infections. Grossly and microscopically, transplanted perichondrium and periosteum allografts appeared to be vital at all postsurgical time-points (Fig. 1, 2) without any signs of allograft rejection. The use of donors with ubiquitous EGFP expression (Lew-Tg(CAG-EGFP)YsRrrc) and EGFP-negative recipients enabled tracing of transplanted cells and their progenies at all studied postsurgical time-points by GFP immunofluorescence (Fig. 3a, 3c).

**Figure 1.**
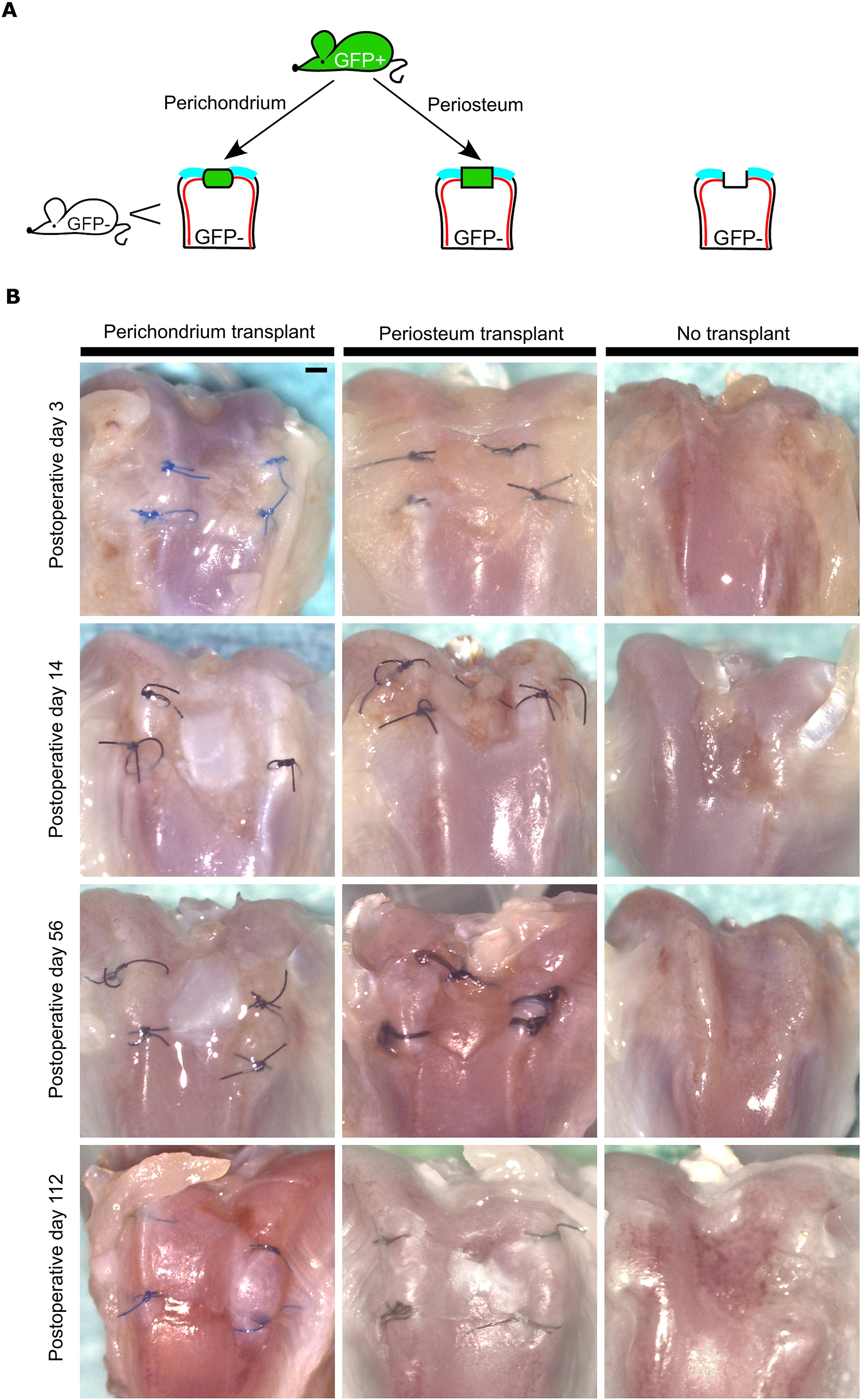
Experimental design and articular surfaces of cartilage defects repaired with perichondrium or periosteum transplants or left untreated. Perichondrium and periosteum grafts were harvested from EGFP positive Lewis (inbred) rats and transplanted into full-thickness cartilage injuries at the distal femoral intercondylar groove in wild-type littermates **(a)**. At post-surgery day 3, 14, 56, and 112, femoral epiphyses were collected, fixed overnight in formalin, decalcified in (EDTA, 15%). Low power microphotographs were captured using a Leica M320 dental microscope **(b)**. Scale bar represents 1000 μm.

**Figure 2.**
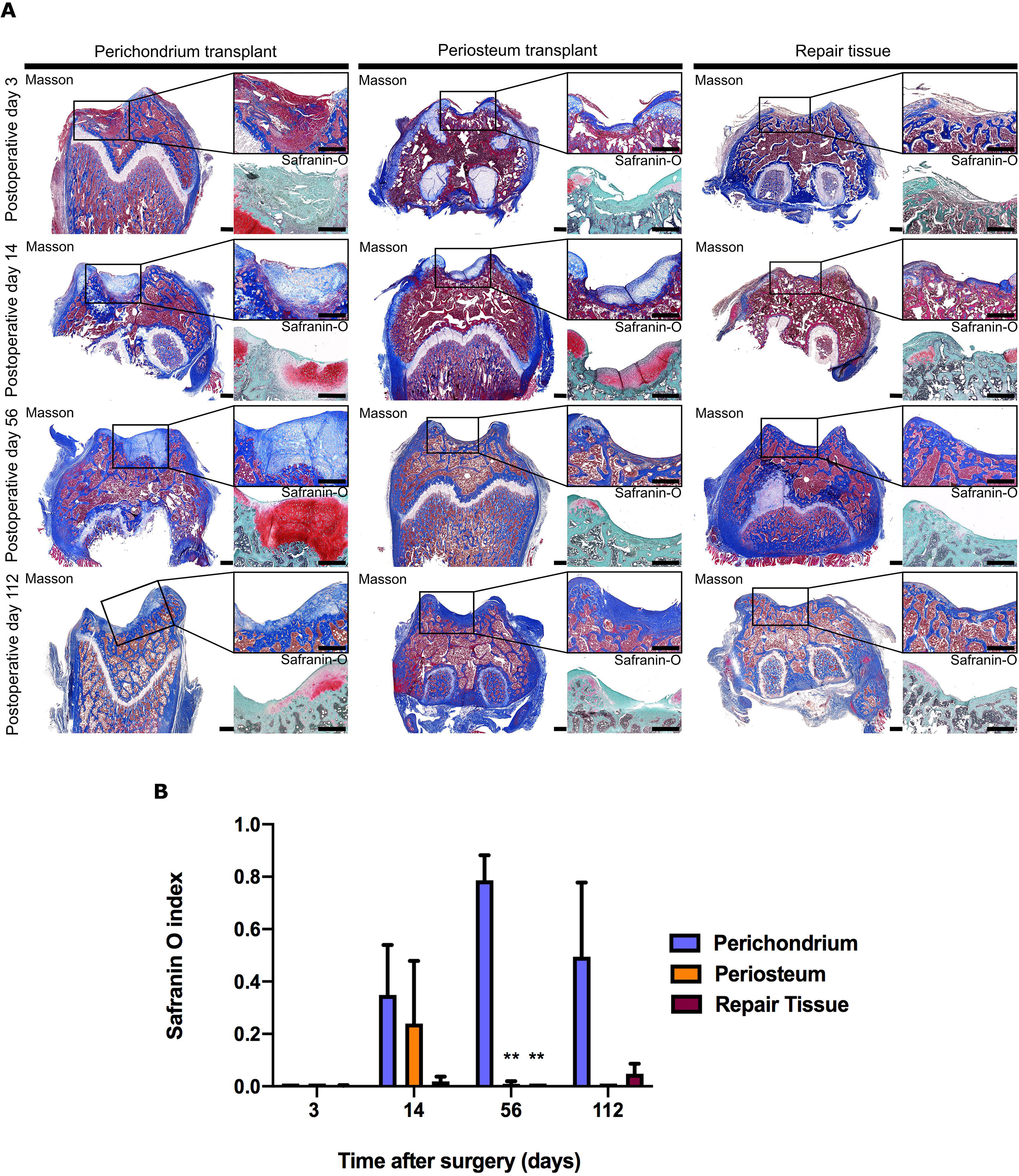
Histology of cartilage defects repaired with perichondrium or periosteum transplants or left untreated. Representative microphotographs of Masson Trichrome and Safranin-O (high power) stained tissue sections of distal femur epiphyses from perichondrium transplanted (n = 3), periosteum transplanted (n = 3) and no transplant control (n = 3 except for day 14 post-surgery (n = 2)) rats at 3, 14, 56, 112 days post-surgery displayed at low (Masson Trichrome) and high (Masson Trichrome and Safranin-O) power magnification **(a)**. Safranin O index was calculated by dividing the safranin O positive area by total area of grafts or repair tissue respectively at 3, 14, 56 and 112 days post-surgery **(b)**. Safranin-O is higher in perichondrium than in periosteum and repair tissue (*P* < 0.001 by ANOVA). (**) compared to perichondrium group; \*\**P* < 0.01. Scale bar represents 500 μm.

**Figure 3.**
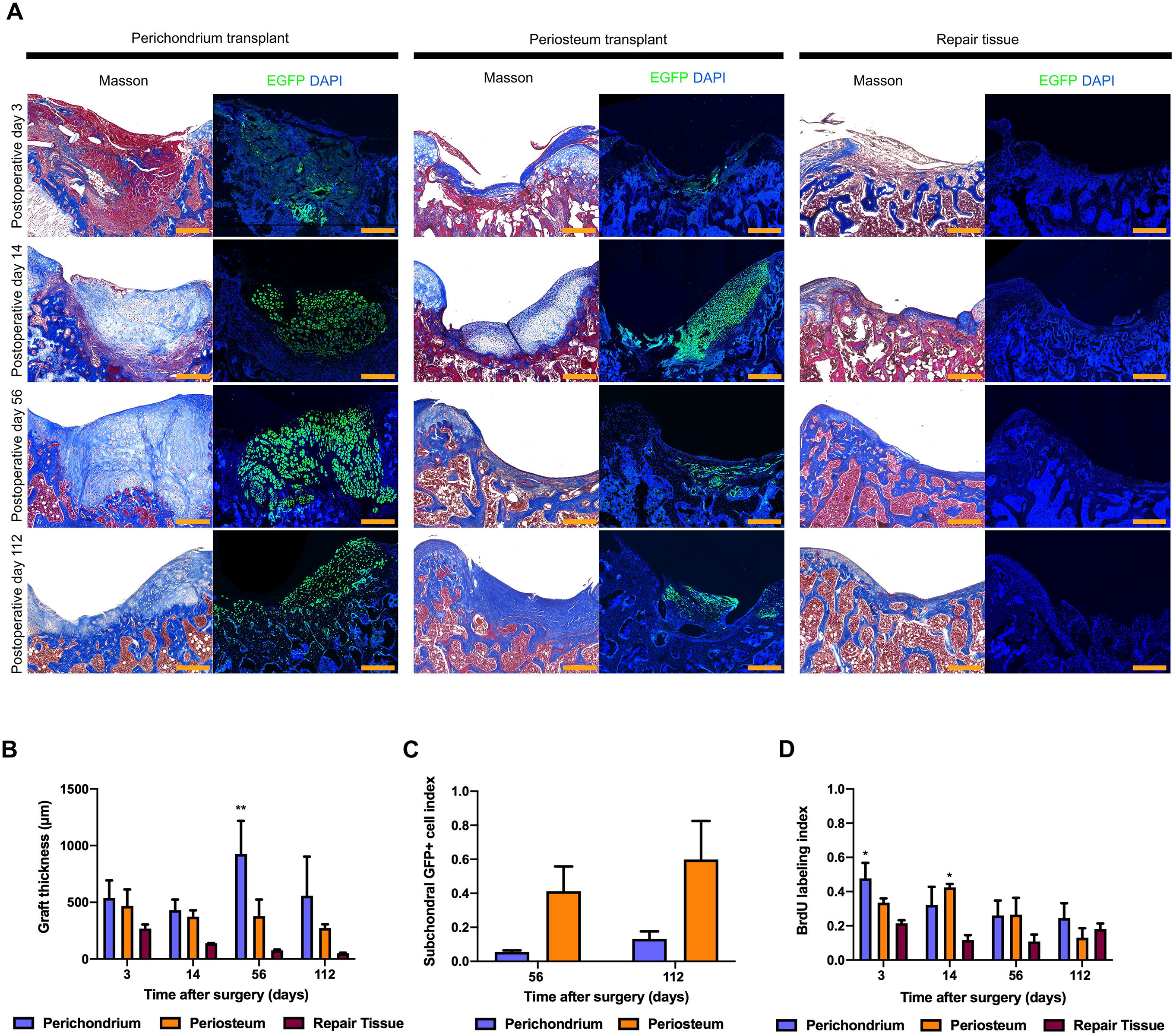
Tracing of transplant derived cells, thickness of grafts and repair tissue, and cell proliferation. Transplant derived cells were visualized using EGFP immunofluorescence on frozen sections of distal epiphyses at post-surgery day 3, 14, 56 and 112. Representative immunofluorescence images were aligned with corresponding Masson’s trichrome stained tissue sections **(a)**. Thickness of grafts and repair tissue **(b)**. Thickness was measured perpendicular to the joint surface at 10 different locations spreading evenly over each graft/repair tissue and averaged. Perichondrium transplants were thicker than periosteum transplants and repair tissue (*P* < 0.001 by ANOVA). Subchondral GFP positive cell index was calculated by dividing the number of positive cells by the total number of cells in the subchondral bone cell compartment **(c)**, and it was higher in periosteum group than in perichondrium group (*P* < 0.05 by ANOVA). Proliferation rate in each transplant and repair tissue was assessed by BrdU labeling and immunohistochemistry. The number of BrdU labelled cells within the graft or repair tissue was divided by the total number of cells to produce BrdU indices **(d)**. Cell proliferation declined with time (*P* < 0.05 by ANOVA). (*) compared to repair tissue group; \**P* < 0.05, \*\**P* < 0.01. Scale bar represents 500 μm.

### Perichondrium transplants underwent robust chondrogenesis and differentiated into hyaline cartilage that expanded and were maintained at post-surgery day 112

At post-surgery day 3, perichondrium grafts were fibrous without any cartilaginous areas detectable on Masson’s trichrome or Safranin O stained slides (Fig. 2) and did not express *Col2a1* (Fig. 4c, 4e), but had high SOX9 expression (Fig. 4a, 4b). In contrast, at 14 days post-surgery the transplanted GFP positive tissue (Fig. 3) consisted almost entirely of light aniline blue and safranin O (Fig. 2) positive cartilaginous tissue and was positive to SOX9 and *Col2a1* at the center of the transplant (Fig. 4a, 4e). This hyaline cartilage phenotype was even more clear at 56 days post-surgery with almost all cells of the GFP positive transplant expressing *Col2a1* and these high expression levels were largely maintained at post-surgery day 112 (Fig. 3a, 4c, 4e). Conversely, perichondrium transplants expressed Col1a1 intensely in the fibrous layer sutured to the subchondral bone at post-surgery day 3 but were mostly negative in the rest of the transplant and at all subsequent time-points (Fig. 4d, 4e).

**Figure 4.**
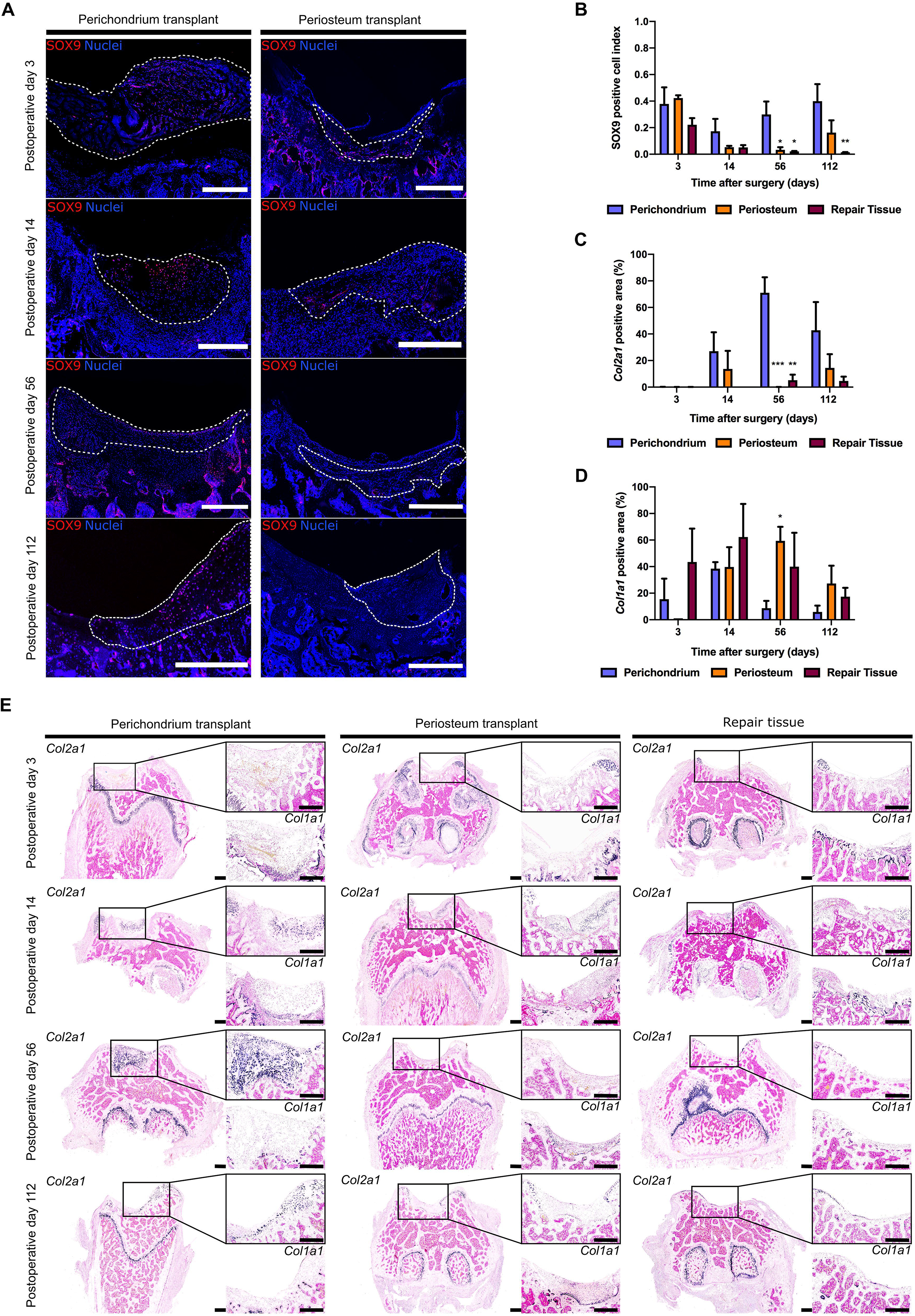
SOX9 immunofluorescence and collagen type I and II *in situ* hybridization of perichondrium and periosteum grafts and repair tissue. SOX9 was targeted by immunofluorescence or immunohistochemistry in frozen or paraffin sections of distal epiphyses at post-surgery day 3, 14, 56 and 112. Representative immunofluorescence images of perichondrium and periosteum transplant groups were displayed with grafts indicated by white dotted lines **(a)**. SOX9 positive cell index was calculated by dividing the number of SOX9 positive cells by total number of cells within grafts or repair tissue at 3, 14, 56, and 112 days post-surgery **(b)**, and was similar in all groups at post-surgery day 3, but then decreased in the periosteum transplant and no transplant groups (*P* < 0.01 by ANOVA). *Col2a1* and *Col1a1* mRNA were visualized (purple coloration) with non-radioactive digoxigenin labeled riboprobes in consecutive paraffin sections of distal femoral epiphyses. *Col2a1* and *Col1a1* positive area percentage was calculated by dividing their positive areas by total area of transplants or repair tissue respectively and multiplied by 100 **(c, d)**. All samples were negative to *Col2a1* at post-surgery day 3, increased to day 14 in both perichondrium and periosteum, and then remained high in perichondrium at post-surgery day 56 and 112, but were mostly negative in periosteum transplants (**c**; *P* < 0.05 and *P* < 0.001 for time and tissue, respectively, by ANOVA). Perichondrium transplants were positive to *Col1a1* in situ hybridization at the bottom of the transplants (fibrous layer of perichondrium) at post-surgery day 3 and 14 but then mostly negative at post-surgery day 56 and 112, whereas periosteum had a high level of Col1a1 expression at the later time-points **(d, e)**. Representative *in situ* hybridization images of perichondrium transplanted, periosteum transplanted and no transplant control groups were displayed in low *(Col2a1)* and high power *(Col2a1 (upper insert) & Col1a1 (lower insert))* magnification (e). (*) Compared to perichondrium transplant group; \**P* < 0.05; \*\**P* < 0.01, \*\*\**P* < 0.001. Scale bar represents 500 μm.

The thickness of perichondrium and periosteum as well as repair tissues and native articular cartilage all declined with time (*P* < 0.01 by ANOVA) with one exception (Fig. 3b). Perichondrium transplant thicknesses increased from day 14 to 56. This increase was due to an early burst in proliferation (Fig.3d, Supplemental Fig. 1a) followed by hypertrophy. At 14 days post-surgery cells with a hypertrophic appearance and positive to *Col10a1* were detected in the central area of the perichondrium transplants (Fig. 2, 5) and also at the bottom of the transplants at post-surgery day 56 and 112 (Fig. 5). Interestingly, at post-surgery day 56 and 112 there were histological signs of active bone remodeling of the hypertrophic cartilage at the bottom of the transplants (Fig. 2a) and there were also GFP positive cells within the bone and bone marrow underneath the transplants (Fig. 3a, 3c). Taken together, these observations demonstrate that transplanted perichondrial cells differentiate into chondrocytes that differentiate into hypertrophic chondrocytes that induce remodeling into bone at the bottom of the transplants. Moreover, GFP positive cells were detected in the subchondral bone suggesting that skeletal progenitor cells of both perichondrium and periosteum are capable of osteoblast and osteocyte differentiation. Interestingly, no hypertrophic differentiation was detected close to the articular surface (Fig. 2, 5) where chondrocytes, instead, remained small, safranin O negative (Fig. 2a) and eventually oriented tangentially to the surface and expressed *Prg4* (Fig. 5b).

**Figure 5.**
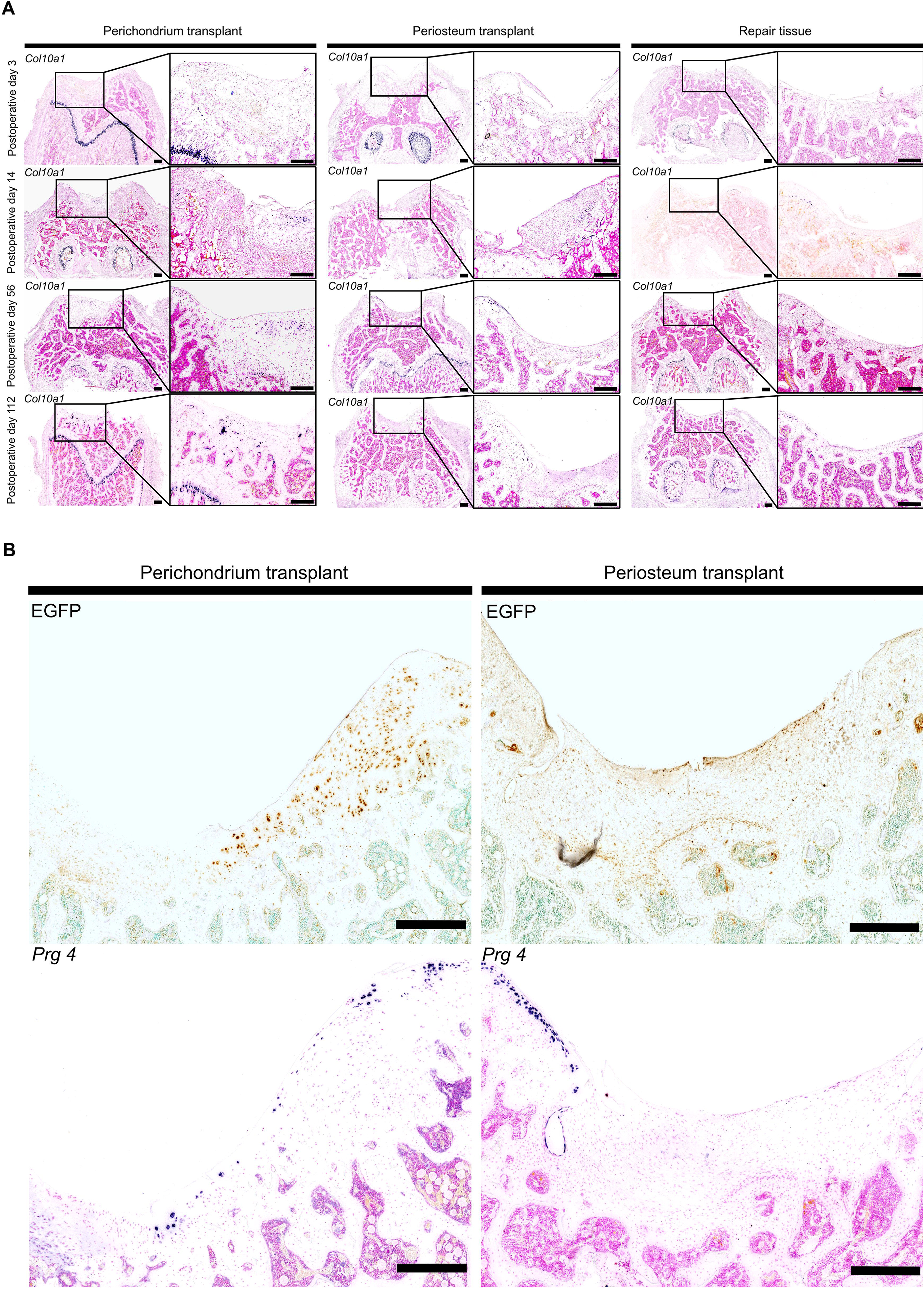
*Col10a1* and *Prg4 in situ* hybridization of perichondrium and periosteum grafts and repair tissue. Representative *Col10a1 in situ* hybridization microphotographs of perichondrium transplanted, periosteum transplanted and no transplant control groups at different post-surgery time-points displayed at low and high power magnification **(a)**. Positive *Col10a1* signal (purple coloration) was visualized at the center of perichondrium transplants at post-surgery day 14, at the center and bottom of the transplants at day 56, and only at the bottom of the perichondrium transplants at post-surgery day 112. GFP immunohistochemistry and *Prg4 in situ* hybridization was performed on consecutive sections to allow for indirect co-localization of GFP protein and *Prg4* mRNA here shown in high power to visualize *Prg4* expression in the most superficial cell layers of perichondrium, but not periosteum, grafts at post-surgery day 112 **(b)**. Scale bar represents 500 μm.

### Periosteum transplants developed into a fibrocartilage structure that was thinning with time

Even though SOX9 expression in periosteum is much lower than in perichondrium at day 0 (Supplemental Fig. 2b, 2c), similar to perichondrium transplants, high numbers of SOX9 positive cells (Fig. 4a, 4b) were observed in periosteum transplants at post-surgery day 3. But it still appeared fibrous with no aniline blue stained cartilage matrix and no Safranin O (Fig. 2a, 2b; *P* < 0.001 by ANOVA) or *Col2a1* positive areas in periosteum transplants at this time-point (Fig. 4c (*P* < 0.001 by ANOVA), 4e). By post-surgery day 14 periosteum transplants had also attained safranin O positive areas (Fig 2a, 2b; *P* < 0.001 by ANOVA) and were also positive for *Col2a1* at the center of the transplants (Fig. 4c, 4e). However, they still had a more fibrocartilage-like appearance and were also positive for *Col1a1* (Fig. 4d, 4e) and had less SOX9 expressing cells (Fig. 4a, 4b). In contrast to the perichondrium transplants, periosteum transplant thickness continuously declined (Fig. 3b; *P* < 0.01 by ANOVA) and lost some of their cartilaginous appearance and cartilage marker expression, i.e. decreased expression of *Col2a1* (Fig. 4c (*P* < 0.01 by ANOVA), 4e) and Safranin O staining (Fig 2a, 2b; *P* < 0.001 by ANOVA). In addition, Col1a1 expression could be detected at all time-points while mostly absent in the transplanted perichondrium (Fig. 4d, 4e). Therefore, injuries repaired with periosteum were thinner (Fig. 3b; *P* = 0.0015 by ANOVA) and had a less cartilaginous appearance than those repaired with perichondrium (Fig. 2, 4). Periosteum transplants were however thicker and had a more cartilaginous appearance than the repair tissue of no transplant controls at the later time-points (Fig. 2, 4). Interestingly, GFP positive periosteum cells did not differentiate into Prg4 expressing chondrocytes at the articular surface (Fig. 5b), and were to a larger extent incorporated into the underlying bone than cells derived from perichondrium transplants (Fig. 3a, 3c (*P* < 0.05 by ANOVA)).

### Cartilage injuries not covered with a transplant healed quickly and provided similar short term, but poor long-term results

Defects left without a transplant were completely filled by fibrous repair tissue already at postoperative day 3 with a macroscopic appearance at least as good as the perichondrium and periosteum filled defects (Fig. 1). Interestingly, the upper part of the tissue filling the defect appeared to be a fibrous sheet continuous with the synovial membrane on one side, therefore giving the appearance of synovial membrane invasion into the injury site (Fig. 1, 2), whereas at the bottom, there were no sharp border to the underlying bone marrow (Fig. 2). Despite having similar levels of SOX9 positive cells at post-surgery day 3 (Fig. 4a, 4b, Supplemental Fig. 1b), repair tissue never attained a hyaline cartilage appearance and remained positive for *Col1a1* by *in situ* hybridization and negative to safranin O staining, as well as for *Col2a1* and *Col10a1* at post-surgery day 14 (Fig. 2, 4, 5) and also at all subsequent time-points. In contrast to perichondrium, but similar to periosteum, the repair tissue thickness declined continuously from post-surgery day 3 to 112 (Fig. 3b; *P* = 0.0015 by ANOVA) at which point there was only a thin layer of translucent fibrocartilage covering the subchondral bone (Fig. 1-3). The repair tissue was negative for *Prg4* expression at 3- and 14-days post-surgery, but an increasing number of *Prg4* positive cells were detected at the surface on post-surgery days 56 and 112 (Supplemental Fig. 2a), suggesting possible infiltration of *Prg4* positive cells from the surrounding articular cartilage and/or synovial tissue.

## DISCUSSION

There is a lack of suitable tissues that can be used to resurface injured and eroded articular cartilage. Rib perichondrium has been used clinically as a tissue source, sometimes with favorable long-term results ^(36,42)^. However, the quality of the resulting joint surfaces and the contribution of the transplanted tissues have not been investigated in detail. We here used transgenic rats with ubiquitous EGFP expression allowing for tracing of transplanted cells. We found that perichondrium transplants formed durable hyaline cartilage tissue with high proteoglycan content that filled out the articular cartilage defects including the area of active remodeling in the subchondral bone, and with time attained structure and chondrocyte marker expression patterns similar to the surrounding articular cartilage. This finding is consistent with the favorable results sometimes reported using perichondrium to resurface injured joints ^(36,42,43)^ and taken together indicate that rib perichondrium is a suitable tissue to repair injured joints.

The levels of SOX9 expression was low in the harvested periosteum compared to perichondrium samples but was then dramatically induced to reach level similar to perichondrium by post-surgery day 3. The increase of SOX9 was sufficient to induce a transient chondrogenic differentiation program of osteochondral progenitors in the periosteum transplants with large parts of the periosteum cells being *Col2a1* positive and surrounded by matrix with high proteoglycan content 14 days after the surgeries. However, in contrast to the perichondrium, the transplanted periosteum tissues expressed *Col1a1* at all time-points and were continuously thinning and lost SOX9 expressing cells and subsequent *Col2a1* expression and proteoglycan content after post-surgery day 14 time-point. Consequently, at post-surgery day 56 and 112 the injuries repaired with periosteum transplants had depressed articular surfaces and an overall histological appearance and chondrocyte marker expression pattern in between the no transplant controls and the perichondrium transplants. The early increase of SOX9 expression was thus not sufficient to induce a lasting chondrogenic program in the periosteum transplants and as the SOX9 expression declined, the cells lost their chondrogenic phenotype and instead differentiated towards osteoblasts. These findings confirm and extend previous studies suggesting that the osteochondroprogenitor cells of periosteum are less chondrogenic and more osteogenic than perichondrial osteochondroprogenitors when placed ectopically in the synovial microenvironment ^(44-46)^. This is also consistent with studies that have demonstrated mostly poor outcomes when periosteum is used as a tissue source for articular cartilage repair ^(47,48)^. The findings are also largely compatible with the physiological role of the periosteum during appositional growth of bones as well as during fracture repair ^(49)^.

A chondrogenic program was induced in perichondrium and partially also in periosteum placed ectopically at the joint surface. This also occurs if perichondrium is placed in the subcutaneous space ^(50,51)^, but not when perichondrium is ectopically placed on muscle or in the liver unless it is exposed to blood ^(30,44,52)^. Our findings taken together with these previous findings may suggest that the synovial microenvironment contains one or several factors that act on perichondrial cells to induce chondrogenic differentiation. In addition, we found that hypertrophic differentiation occurred first at the center and later at the bottom of the transplant, but never at the surface, and also that *Prg4* positive cells were detected at the surface of the formed cartilage at the latest time-point. This observation might suggest that the cartilage formed from the perichondrium transplant has an inherent chondrogenic program producing a cartilaginous structure with chondrocyte differentiation marker expression profile similar to articular cartilage. However, a more likely explanation would be that the perichondrium has an inherent chondrogenic program that is modified by the local microenvironment of the joint. Specifically, that the joint microenvironment may inhibit hypertrophic differentiation and promote differentiation towards the superficial layer chondrocyte phenotype ^(53,54)^. This hypothesis would explain our observation that the newly formed chondrocytes of the perichondrium underwent hypertrophic differentiation only some distance away from the surface, but never close to the surface where chondrocytes instead remained small and eventually attained a superficial chondrocyte-like phenotype.

Articular cartilage injuries left without a transplant were quickly filled by fibrous tissue that appeared to restore joint function but never differentiated into hyaline cartilage. This is consistent with the current thinking that full thickness cartilage injuries are repaired presumably by a blood clot from the bone marrow, which is organized into a fibrocartilage tissues that, in the long-term, leave the injured joints susceptible to future degeneration and osteoarthritis development. However, macroscopic and microscopic observations in our study suggest that in addition to the formation of a blood clot, rapid synovial invasion covering the blood clot and the injury also occurs. However, there were almost no *Prg4* positive cells at the injury site at post-surgery day 3 and 14. This could be due to a loss of *Prg4* expression in synovial cells as they were attracted to the injury site, or that only *Prg4* negative synovial tissue cells were attracted to the injury site. Other options can be that the injury site was mostly filled-up by bone marrow cells and that the blood clot did not only fill up the injury but also became continuous with the synovial membrane during the early healing phase. Further studies are needed to address these possibilities as we were not able to definitely distinguish between these options in the current study. It is interesting that a rapid and efficient repair of cartilage injuries has evolved, but the repair tissue never forms a durable hyaline cartilage that fills the injury, forming instead only a thinning layer of fibrocartilage that leaves the injured joint susceptible to osteoarthritis development. This may be due to the lack of suitable skeletal stem cells with chondrogenic potential and/or that the local microenvironment prevents chondrogenic differentiation of available stem cells. A recent study indicates that this limitation of the endogenous repair tissue may be overcome if the synovial microenvironment is modified. In the study, Murphy et al showed that co-delivery of BMP2 and a soluble VEGFR1 in knee joints at microfracture surgery was sufficient to induce SOX9 expression and formation of hyaline cartilage in the repair tissue ^(55)^.

Free, autologous perichondrium transplants have been used successfully, but with variable results, to repair a variety of articular cartilage injuries ^(32-35,56,57)^. We used a rat model allowing for tracing of transplanted cells and found that surfaces repaired with perichondrium were exclusively made up of transplanted cells and their progenies even after 112 days, thus addressing the long-standing question whether perichondrium itself have the capacity to regenerate a new joint surface or if it merely attracts invasion of chondrogenic stem cells from the bone marrow, synovial membrane or other surrounding tissues ^(21)^, which in turn regenerate the surface. The finding that the transplanted perichondrium expanded and filled the articular cartilage defects and that the surfaces seemed to differentiate and mold to form an articular cartilage-like covering of the subchondral bone, is consistent with clinical observations that perichondrium transplantation in many cases can produce favorable long-term results if transplant detachment and other potential short-term complications are avoided ^(36,42)^.

In summary, rib perichondrium transplanted to full-thickness articular cartilage defects produced hyaline cartilage that filled out the defects and with time differentiated and molded to attain a structure and chondrocyte marker expression pattern similar to the surrounding articular cartilage. In contrast, transplanted periosteum resulted in a thinning layer of fibrocartilage covering a crater-like defect that with time provided more cells into the underlying bone than the articular cartilage itself. These findings indicate that perichondrium, but not periosteum, is a suitable tissue-source for repair of articular cartilage defects, resurfacing of injured joints, and tissue engineering of articular cartilage. In addition, we suggest that the most likely explanation for the absence of hypertrophic differentiation in the superficial cell layers and their differentiation towards superficial-like cells is that the synovial microenvironment inhibits chondrocyte hypertrophy and promote articular cartilage differentiation. Further studies exploring the role of the synovial joint microenvironment as well as the possibilities of using perichondrium as a tissue source for articular cartilage repair, resurfacing, and articular cartilage tissue engineering are warranted.

## Acknowledgements

The authors acknowledge expert advice from Dr. Phillip Newton in Department of Women’s and Children’s Health at Karolinska Institutet for Safranin-O staining and for immunofluorescence confocal microscopy. This study was supported by grants from the Swedish Research Council (project K2015–54X-22 736–01–4 & 2015-02227), the Swedish Governmental Agency for Innovation Systems (Vinnova) (2014-01438), Marianne and Marcus Wallenberg Foundation, the Stockholm County Council, the Uppsala County Council, Byggmästare Olle Engkvist Stiftelse, the Swedish Society of Medicine, Novo Nordisk Foundation, Erik och Edith Fernström Foundation for Medical Research, HKH Kronprinsessan Lovisas förening för barnasjukvård, Sällskapet Barnavård, Stiftelsen Frimurare Barnhuset i Stockholm, Promobilia, Nyckelfonden, and Karolinska Institutet, Stockholm, Sweden, and Örebro University, Örebro, Sweden.

## Author contributions

O.N. and T.V. conceived the project and designed the study. Z.D., D.M., M.B., A.B., A.G., T.V. L.O., and O.N. performed experiments, O.N., T.V., Z.D., and D.M. analyzed data and wrote the manuscript. All authors read and approved the final draft.

---

**Supplemental Figure 1. Cell proliferation and SOX9 immunofluorescence in repair tissue group** Cell proliferation was assessed by BrdU labeling followed by BrdU immunohistochemistry on paraffin sections of distal epiphyses from grafts and repair tissues at post-surgery day 3, 14, 56 and 112. Representative immunohistochemistry high power magnification images were aligned with lower power magnification Masson’s trichrome staining images from consecutive sections **(a)**. SOX9 expression was visualized by immunofluorescence on frozen sections of distal femoral epiphyses. Representative SOX9 immunofluorescence images of repair tissue at post-surgery day 3, 14, 56 and 112 **(b)**. Scale bar represents 500 μm.

**Supplemental Figure 2. *Prg4* in situ hybridization of perichondrium, periosteum grafts and repair tissue at post-surgery and markers expression in grafts at day 0.** Representative *Prg4* in situ hybridization images of perichondrium transplanted, periosteum transplanted and no transplant control groups displayed at low and high power magnification **(a)**. The relative mRNA expression of *Col2a1, Col10a1, SOX9, ACAN*, and *Prg4* in perichondrium and periosteum pieces (3 technical replicates performed in each group with tissue pieces from same donor animal) assessed by real-time PCR at day 0 **(b)**. Representative Masson’s trichrome staining and SOX9 immunohistochemistry images of perichondrium and periosteum transplant sections at day 0 **(c)**. Scale bar represents 200 μm in low and 20 μm in high magnification images.

## References

1. Kronenberg HM. Developmental regulation of the growth plate. Nature. May 15 2003;423(6937):332–6. Epub 2003/05/16.

2. Gerber HP, Vu TH, Ryan AM, Kowalski J, Werb Z, Ferrara N. VEGF couples hypertrophic cartilage remodeling, ossification and angiogenesis during endochondral bone formation. Nat Med. Jun 1999;5(6):623–8. Epub 1999/06/17.

3. Patel JM, Dunn MG. Cartilage tissue engineering. Regenerative Engineering of Musculoskeletal Tissues and Interfaces2015. p. 135–60.

4. Ono N, Balani DH, Kronenberg HM. Stem and progenitor cells in skeletal development. Curr Top Dev Biol. 2019;133:1–24. Epub 2019/03/25.

5. Kronenberg HM. The role of the perichondrium in fetal bone development. Ann N Y Acad Sci. Nov 2007;1116:59–64. Epub 2007/12/18.

6. Ono N, Kronenberg HM. Bone repair and stem cells. Curr Opin Genet Dev. Oct 2016;40:103–7. Epub 2016/07/12.

7. Colnot C. Skeletal cell fate decisions within periosteum and bone marrow during bone regeneration. J Bone Miner Res. Feb 2009;24(2):274–82. Epub 2008/10/14.

8. Lefebvre V, Bhattaram P. Vertebrate skeletogenesis. Curr Top Dev Biol. 2010;90:291–317. Epub 2010/08/10.

9. Shwartz Y, Viukov S, Krief S, Zelzer E. Joint Development Involves a Continuous Influx of Gdf5-Positive Cells. Cell Rep. Jun 21 2016;15(12):2577–87. Epub 2016/06/14.

10. Khan IM, Redman SN, Williams R, Dowthwaite GP, Oldfield SF, Archer CW. The development of synovial joints. Curr Top Dev Biol. 2007;79:1–36. Epub 2007/05/15.

11. Hyde G, Boot-Handford RP, Wallis GA. Col2a1 lineage tracing reveals that the meniscus of the knee joint has a complex cellular origin. J Anat. Nov 2008;213(5):531–8. Epub 2008/11/19.

12. Koyama E, Shibukawa Y, Nagayama M, Sugito H, Young B, Yuasa T, et al. A distinct cohort of progenitor cells participates in synovial joint and articular cartilage formation during mouse limb skeletogenesis. Dev Biol. Apr 1 2008;316(1):62–73. Epub 2008/02/26.

13. Pacifici M, Koyama E, Iwamoto M. Mechanisms of synovial joint and articular cartilage formation: recent advances, but many lingering mysteries. Birth Defects Res C Embryo Today. Sep 2005;75(3):237–48. Epub 2005/09/28.

14. Kozhemyakina E, Lassar AB, Zelzer E. A pathway to bone: signaling molecules and transcription factors involved in chondrocyte development and maturation. Development. Mar 1 2015;142(5):817–31. Epub 2015/02/26.

15. Li L, Newton PT, Bouderlique T, Sejnohova M, Zikmund T, Kozhemyakina E, et al. Superficial cells are self-renewing chondrocyte progenitors, which form the articular cartilage in juvenile mice. FASEB J. Mar 2017;31(3):1067–84. Epub 2016/12/15.

16. Chijimatsu R, Saito T. Mechanisms of synovial joint and articular cartilage development. Cell Mol Life Sci. Oct 2019;76(20):3939–52. Epub 2019/06/16.

17. Buckwalter JA. Were the Hunter brothers wrong? Can surgical treatment repair articular cartilage? Iowa Orthop J. 1997;17:1–13. Epub 1997/01/01.

18. Mankin HJ. The response of articular cartilage to mechanical injury. J Bone Joint Surg Am. Mar 1982;64(3):460–6. Epub 1982/03/01.

19. Hunziker EB, Lippuner K, Keel MJ, Shintani N. An educational review of cartilage repair: precepts & practice--myths & misconceptions--progress & prospects. Osteoarthritis Cartilage. Mar 2015;23(3):334–50. Epub 2014/12/24.

20. Hunziker EB, Rosenberg LC. Repair of partial-thickness defects in articular cartilage: cell recruitment from the synovial membrane. J Bone Joint Surg Am. May 1996;78(5):721–33. Epub 1996/05/01.

21. Hunziker EB. Articular cartilage repair: basic science and clinical progress. A review of the current status and prospects. Osteoarthritis Cartilage. Jun 2002;10(6):432–63. Epub 2002/06/12.

22. Lynch TS, Patel RM, Benedick A, Amin NH, Jones MH, Miniaci A. Systematic review of autogenous osteochondral transplant outcomes. Arthroscopy. Apr 2015;31(4):746–54. Epub 2015/01/27.

23. Angele P, Niemeyer P, Steinwachs M, Filardo G, Gomoll AH, Kon E, et al. Chondral and osteochondral operative treatment in early osteoarthritis. Knee Surg Sports Traumatol Arthrosc. Jun 2016;24(6):1743–52. Epub 2016/02/29.

24. Campbell AB, Pineda M, Harris JD, Flanigan DC. Return to Sport After Articular Cartilage Repair in Athletes’ Knees: A Systematic Review. Arthroscopy. Apr 2016;32(4):651–68 e1. Epub 2015/11/04.

25. Brittberg M, Lindahl A, Nilsson A, Ohlsson C, Isaksson O, Peterson L. Treatment of deep cartilage defects in the knee with autologous chondrocyte transplantation. N Engl J Med. Oct 6 1994;331(14):889–95. Epub 1994/10/06.

26. O’Driscoll SW, Fitzsimmons JS. The role of periosteum in cartilage repair. Clin Orthop Relat Res. Oct 2001(391 Suppl):S190–207. Epub 2001/10/18.

27. Skoog T, Johansson SH. The formation of articular cartilage from free perichondrial grafts. Plast Reconstr Surg. Jan 1976;57(1):1–6. Epub 1976/01/01.

28. Ohlsen L, Skoog T, Sohn SA. The pathogenesis of cauliflower ear. An experimental study in rabbits. Scand J Plast Reconstr Surg. 1975;9(1):34–9. Epub 1975/01/01.

29. Engkvist O, Johansson SH, Ohlsen L, Skoog T. Reconstruction of articular cartilage using autologous perichondrial grafts. A preliminary report. Scand J Plast Reconstr Surg. 1975;9(3):203–6. Epub 1975/01/01.

30. Ohlsen L. Cartilage formation from free perichondrial grafts: an experimental study in rabbits. Br J Plast Surg. Jul 1976;29(3):262–7. Epub 1976/07/01.

31. Engkvist O, Ohlsen L. Reconstruction of articular cartilage with free autologous perichondrial grafts. An experimental study in rabbits. Scand J Plast Reconstr Surg. 1979;13(2):269–74. Epub 1979/01/01.

32. Engkvist O, Johansson SH. Perichondrial arthroplasty. A clinical study in twenty-six patients. Scand J Plast Reconstr Surg. 1980;14(1):71–87. Epub 1980/01/01.

33. Seradge H, Kutz JA, Kleinert HE, Lister GD, Wolff TW, Atasoy E. Perichondrial resurfacing arthroplasty in the hand. J Hand Surg Am. Nov 1984;9(6):880–6. Epub 1984/11/01.

34. Pastacaldi P. Perichondrial wrist arthroplasty--a follow-up study in 17 rheumatoid patients. Ann Plast Surg. Aug 1982;9(2):146–51. Epub 1982/08/01.

35. Katsaros J, Milner R, Marshall NJ. Perichondrial arthroplasty incorporating costal cartilage. J Hand Surg Br. Apr 1995;20(2):137–42. Epub 1995/04/01.

36. Muder D, Nilsson O, Vedung T. Reconstruction of finger joints using autologous rib perichondrium - an observational study at a single Centre with a median follow-up of 37 years. BMC Musculoskelet Disord. Apr 29 2020;21(1):278. Epub 2020/05/01.

37. Nilsson O, Parker EA, Hegde A, Chau M, Barnes KM, Baron J. Gradients in bone morphogenetic protein-related gene expression across the growth plate. J Endocrinol. Apr 2007;193(1):75–84. Epub 2007/04/03.

38. Chau M, Lui JC, Landman EB, Spath SS, Vortkamp A, Baron J, et al. Gene expression profiling reveals similarities between the spatial architectures of postnatal articular and growth plate cartilage. PLoS One. 2014;9(7):e103061. Epub 2014/07/30.

39. Spath SS, Andrade AC, Chau M, Baroncelli M, Nilsson O. Evidence That Rat Chondrocytes Can Differentiate Into Perichondrial Cells. JBMR Plus. Nov 2018;2(6):351–61. Epub 2018/11/22.

40. Bandyopadhyay A, Kubilus JK, Crochiere ML, Linsenmayer TF, Tabin CJ. Identification of unique molecular subdomains in the perichondrium and periosteum and their role in regulating gene expression in the underlying chondrocytes. Dev Biol. Sep 1 2008;321(1):162–74. Epub 2008/07/08.

41. Lui JC, Chau M, Chen W, Cheung CS, Hanson J, Rodriguez-Canales J, et al. Spatial regulation of gene expression during growth of articular cartilage in juvenile mice. Pediatr Res. Mar 2015;77(3):406–15. Epub 2014/12/19.

42. Muder D, Hailer NP, Vedung T. Two-component surface replacement implants compared with perichondrium transplantation for restoration of Metacarpophalangeal and proximal Interphalangeal joints: a retrospective cohort study with a mean follow-up time of 6 respectively 26 years. BMC Musculoskelet Disord. Oct 7 2020;21(1):657. Epub 2020/10/09.

43. Bouwmeester PS, Kuijer R, Homminga GN, Bulstra SK, Geesink RG. A retrospective analysis of two independent prospective cartilage repair studies: autogenous perichondrial grafting versus subchondral drilling 10 years post-surgery. J Orthop Res. Mar 2002;20(2):267–73. Epub 2002/04/02.

44. Skoog V, Widenfalk B, Ohlsen L, Wasteson A. The effect of growth factors and synovial fluid on chondrogenesis in perichondrium. Scand J Plast Reconstr Surg Hand Surg. 1990;24(2):89–95. Epub 1990/01/01.

45. Ritsila V, Alhopuro S, Rintala A. Bone formation with free periosteum. An experimental study. Scand J Plast Reconstr Surg. 1972;6(1):51–6. Epub 1972/01/01.

46. Ritsila V. Regeneration of articular cartilage defects with free perichondrial grafts. Clinical Orthopaedics and Related Research. 1994;302:259–65.

47. Gooding CR, Bartlett W, Bentley G, Skinner JA, Carrington R, Flanagan A. A prospective, randomised study comparing two techniques of autologous chondrocyte implantation for osteochondral defects in the knee: Periosteum covered versus type I/III collagen covered. Knee. Jun 2006;13(3):203–10. Epub 2006/04/29.

48. McCarthy HS, Roberts S. A histological comparison of the repair tissue formed when using either Chondrogide((R)) or periosteum during autologous chondrocyte implantation. Osteoarthritis Cartilage. Dec 2013;21(12):2048–57. Epub 2013/10/29.

49. van Gastel N, Stegen S, Eelen G, Schoors S, Carlier A, Daniels VW, et al. Lipid availability determines fate of skeletal progenitor cells via SOX9. Nature. Mar 2020;579(7797):111–7. Epub 2020/02/28.

50. ten Koppel PG, van Osch GJ, Verwoerd CD, Verwoerd-Verhoef HL. Efficacy of perichondrium and a trabecular demineralized bone matrix for generating cartilage. Plast Reconstr Surg. Nov 1998;102(6):2012–20; discussion 21. Epub 1998/11/12.

51. Kagimoto S, Takebe T, Kobayashi S, Yabuki Y, Hori A, Hirotomi K, et al. Autotransplantation of Monkey Ear Perichondrium-Derived Progenitor Cells for Cartilage Reconstruction. Cell Transplant. 2016;25(5):951–62. Epub 2016/02/18.

52. Sari A, Tuncer S, Ayhan S, Elmas C, Ozogul C, Latifoglu O. What wrapped perichondrial and periosteal grafts offer as regenerators of new tissue. J Craniofac Surg. Nov 2006;17(6):1137–43. Epub 2006/11/23.

53. Dou Z. Growth plate cartilage transplanted to the articular surface remodels into articular-like cartilage in a process promoted by the synovial joint microenvironment. Bone Report. 2020;13S:Abstract 100410.

54. Baroncelli M. Synovial cells secrete a temperature-stable protein that inhibits hypertrophic differentiation and induces articular cartilage differentiation of chondrocytes in vitro. Bone Reports. 2020;13S:Abstract 100665.

55. Murphy MP, Koepke LS, Lopez MT, Tong X, Ambrosi TH, Gulati GS, et al. Articular cartilage regeneration by activated skeletal stem cells. Nat Med. Oct 2020;26(10):1583–92. Epub 2020/08/19.

56. Bouwmeester SJ, Beckers JM, Kuijer R, van der Linden AJ, Bulstra SK. Long-term results of rib perichondrial grafts for repair of cartilage defects in the human knee. Int Orthop. 1997;21(5):313–7. Epub 1997/01/01.

57. Vedung T, Vinnars B. Resurfacing the distal radioulnar joint with rib perichondrium-a novel method. J Wrist Surg. Aug 2014;3(3):206–10. Epub 2014/08/07.

