## Supplementary figures and images for "Rat perichondrium transplanted to articular cartilage defects forms articular-like, hyaline cartilage"

### Supplemental figure 1

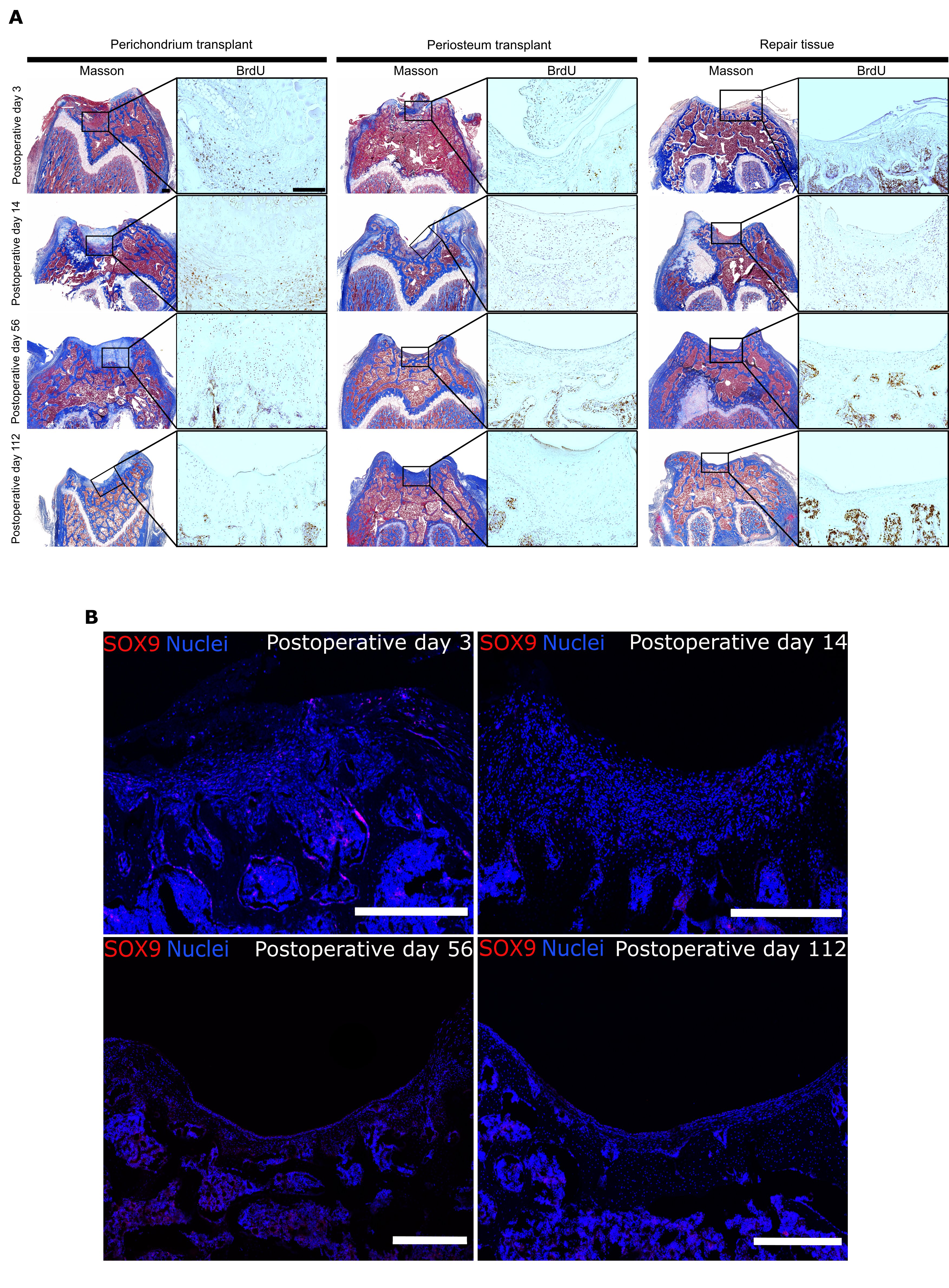

### Supplemental figure 2

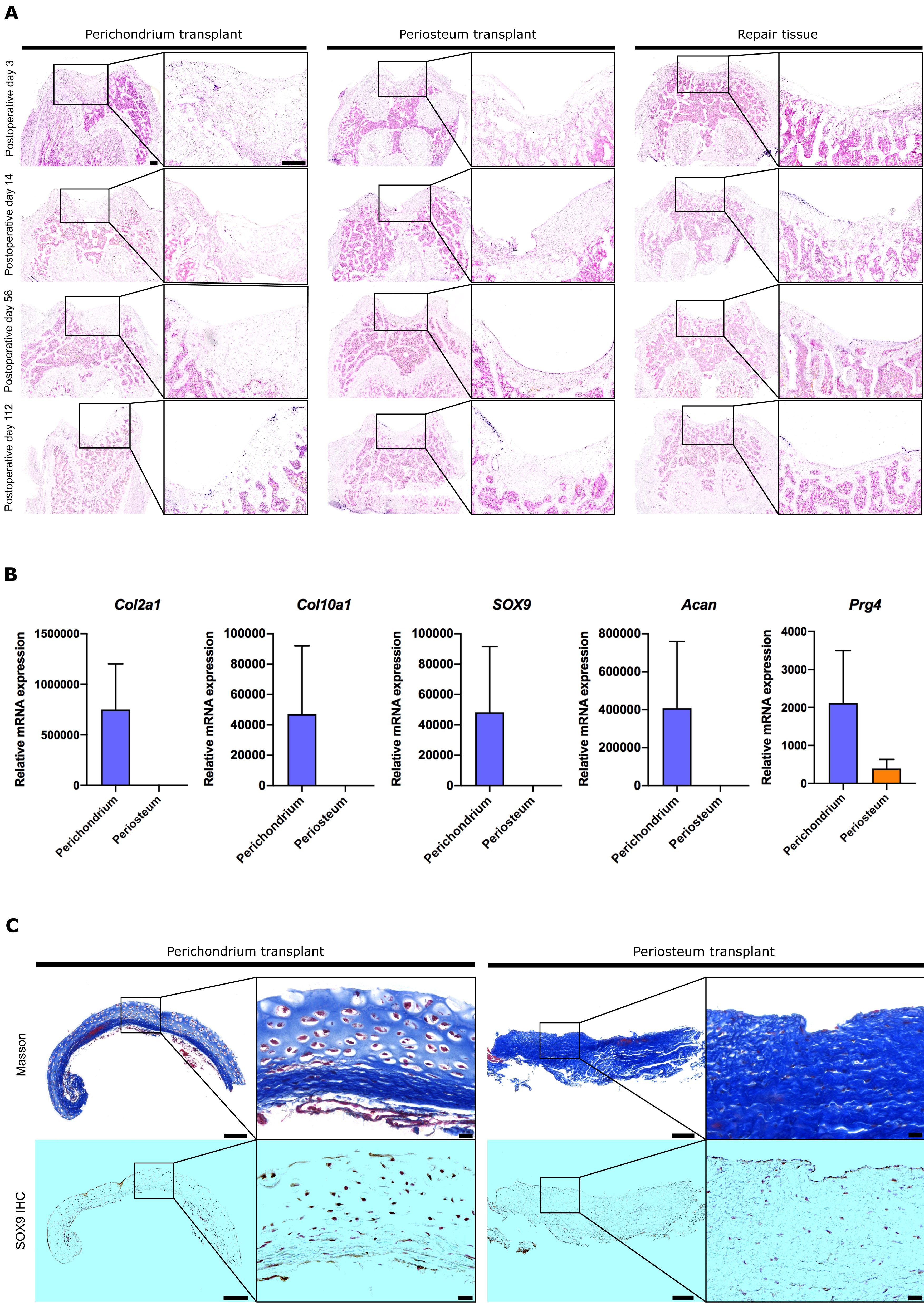
