## Supplemental table 1 for "Rat perichondrium transplanted to articular cartilage defects forms articular-like, hyaline cartilage"

**Supplementary Table 1.** List of primer sets used for qPCR in this study. Genes were quantified using SYBR otherwise indicated.

| ***Prg4*** |  |
| --- | --- |
| Forward primer: | 5' GCATTAACATCCATCCCATGTTT 3' |
| Reverse primer: | 5' CCATCCACTGGCTTACCATTG 3' |
| ***Col2a1*** |  |
| Forward primer: | 5' GCCAGGATGCCCGAAAATTAG 3' |
| Reverse primer: | 5' CCACCAGCCTTCTCGTCAAA 3' |
| ***Acan*** |  |
| Forward primer: | 5' GTGCGCCCATCATCAGAAAC 3' |
| Reverse primer: | 5' GGTGCTTGGACAGTGGATCA 3' |
| ***Col10a1*** |  |
| Taqman | Rn01408030_m1 |
| ***SOX9*** |  |
| Taqman | Rn01751070_m1 |
| ***18s*** |  |
| Taqman | X03205.1 |
